# Visual speech enhances phoneme separability in human superior temporal gyrus

**DOI:** 10.64898/2026.09.10.750762

**Authors:** Yike Li, Iain DeWitt, Jonathan R. Brennan, Vibhangini S. Wasade, William Stacey, David Brang

## Abstract

Visual speech, such as lipreading, facilitates spoken word recognition, but the neural mechanisms underlying audiovisual speech perception remain poorly understood. Visual cues may disambiguate fine-grained articulatory features during early perceptual stages or instead integrate with speech at more categorical, phoneme-level stages. To test how speech representations are modulated by visual input, we analyzed intracranial electroencephalography (iEEG) signals recorded from 12 epilepsy patients performing an audiovisual speech perception task. Participants perceived 16 monosyllabic words presented in auditory-only, visual-only, or congruent audiovisual formats. Words were constructed from four onset consonants (/b/, /g/, /m/, /n/) and four rimes (vowel nucleus and any coda consonants). We examined event-related potentials (ERP) in superior temporal gyrus (STG) and trained support vector machine (SVM) classifiers to decode word identity from neural activity at individual electrodes. Discrete and continuous confusion matrices captured complementary changes in classification accuracy and normalized inverse classification loss, a continuous proxy for classifier confidence. Decoding performance was hierarchically evaluated at the word, phoneme, and phonetic feature levels to determine the representations affected by visual speech. Congruent audiovisual speech increased classifier confidence for phoneme-level representations and improved decoding accuracy at both the word and phoneme level, without corresponding effects on phonetic features. Time-resolved analyses further revealed earlier successful decoding for audiovisual than auditory-only speech, with audiovisual enhancement primarily observed for onset consonants rather than rimes. Together, these findings suggest that visual speech sharpens primarily categorical phoneme representations in STG, with accelerated speech processing and improved word recognition emerging as downstream consequences of phoneme-level enhancement.

**Significance Statement:** Seeing a speaker’s face improves speech perception, especially in noise, but the neural representations altered by visual speech remain unclear. Using intracranial recordings from human superior temporal gyrus, we show that congruent visual speech does not simply amplify all speech-related information. Instead, visual input selectively enhances the separability of confusable phoneme-level representations and improves word identity decoding, while leaving the robust phonetic-feature organization largely intact. These findings clarify the distinction between what visual speech modulates and the representational structure of speech in auditory cortex, suggesting that audiovisual facilitation primarily acts on categorical speech representations that more directly support word recognition.

## Introduction

Speech perception is a special case of multisensory processing (Sumby & Pollack, 1954). Although the normal-hearing population relies predominantly on audition, incongruent visual information reshapes the auditory phoneme percept (Alsius et al., 2018; McGurk & MacDonald, 1976), and congruent cues enhance speech comprehension in optimal and challenging environments (Ross et al., 2007; Sumby & Pollack, 1954). Despite extensive behavioral evidence, *how* visual speech alters auditory cortical processing remains unclear.

Visual speech modulates auditory processing across temporal and spectral scales (Cao et al., 2024; Karthik et al., 2021; Mégevand et al., 2020), and silent lipreading alone activates auditory cortex (Pekkola et al., 2005). Recent intracranial electroencephalography (iEEG) and fMRI results suggest that beyond the modulatory effects on auditory processing, visemes—the categorical units of speech-related lip movements (Fisher, 1968)—may be encoded in auditory cortex (Audenhaege et al., 2025; Karthik et al., 2024). Visual information is thus relayed into auditory cortex, yet its interaction with linguistic representations is underexplored.

Linguistic representations (DeWitt & Rauschecker, 2013; Gwilliams, Bhaya-Grossman, et al., 2025; Hickok & Poeppel, 2004, 2007; Holt & Lotto, 2010) and their organization in superior temporal gyrus (STG) (Bhaya-Grossman et al., 2025; de Heer et al., 2017) recapitulate the hierarchical structure of speech. Speech sounds are described along composable articulatory dimensions, which define phonetic features such as place of articulation (POA, where the vocal tract constricts airflow) and manner of articulation (MOA, how airflow is constricted). Phonemes are language-specific minimal units distinguishing word meanings. In STG, phonetic features are represented by local spectrotemporal tunings (Leonard et al., 2024; Mesgarani et al., 2014), whereas phonemes are categorically organized at the population level (Chang et al., 2010). To achieve stable yet flexible speech perception despite variability (Kleinschmidt & Jaeger, 2015), listeners integrate available cues to resolve ambiguous phonemes or restore phonetic features (Cole, 1973; Gwilliams et al., 2018; Leonard et al., 2016). Computational models further support efficient multisensory cue combination (Chandrasekaran, 2017; Körding et al., 2007; Magnotti & Beauchamp, 2017), raising the question of where in the hierarchy congruent visual cues are integrated.

While all levels of the psycholinguistic hierarchy are susceptible to visual influence (Bernstein & Liebenthal, 2014; Campbell, 2008), visual speech is often thought to enrich phonetic features, the acoustic consequences of articulator states (Grant & Walden, 1996; Summerfield, 1979). Accordingly, facial cues bias perceived POA even when acoustics are identical (Green & Kuhl, 1989), and McGurk-style paradigms show systematic shifts in phonetic encoding (Brancazio et al., 2003; Shahin et al., 2018). Consistent with this feature-based account, neurophysiologically, visual speech produces articulator-specific facilitation of early auditory processing proportional to its phonetic predictiveness (van Wassenhove et al., 2005).

Alternatively, with meaning-rich stimuli, the goal of audiovisual speech is intelligibility (Sumby & Pollack, 1954), *i.e.*, improving phoneme and word perception rather than refining features. Visual speech alone supports word identification (Bernstein et al., 2000), and intracranial evidence shows stronger audiovisual benefits for mouth-leading than voice-leading words (Karas et al., 2019). Visual speech may therefore directly modulate phoneme-level processing by suppressing incompatible alternatives to restrict lexical candidates (Luce, 1986; Tye-Murray et al., 2007). However, as phonetic features underlie phoneme category membership, the two levels are intertwined, and phoneme judgments remain sensitive to acoustic similarity (Samuel, 1981), making it challenging to isolate the representational level at which visual input shapes word perception.

Here we test how visual speech interacts with auditory representations at phonetic-feature, phoneme, and word levels during word identification. We recorded iEEG from STG in epilepsy patients hearing and viewing monosyllabic words and decoded word identity using support vector machines. Decoding was evaluated hierarchically with confusion and confidence matrices, capturing discrete changes in classifier decisions and continuous representational gains, respectively. Congruent visual speech increased phoneme-and word-level decoding accuracy and phoneme-level classifier confidence, without comparable benefits for isolated phonetic features. Visual speech thus appears to enhance the inter-category separability of phoneme representations in human STG, with word-level gains emerging downstream from enhanced phoneme separability.

## Methods

### Experimental model and participant details

#### Participants

All experimental procedures were approved by the Institutional Review Boards (IRBs) at the University of Michigan (UM) and Henry Ford (HF) hospitals. IEEG data were recorded from 13 patients undergoing clinical evaluations for intractable epilepsy. One patient was excluded from the analysis due to a lack of STG coverage, resulting in a final sample of *n =* 12 patients (5 female; mean age = 30.0, *std* = 9.36 years; 8 right-handed, 2 left-handed, and 2 with no clear hand preference). All patients were native speakers of American English and provided written informed consent to participate in the research. Electrode type and implant location were decided solely based on clinical needs. All 12 analyzed patients had sEEG depth electrodes. 11 patients were implanted only with depth electrodes, with 5-mm center-to-center spacing for UM patients and 3.5-mm spacing for HF patients. One patient additionally had a subdural ECoG grid with 10-mm center-to-center spacing implanted above the left STG. Two patients had bilateral STG coverage, whereas the remaining 10 had left STG coverage only. For visualization purposes, all electrodes were projected onto the left hemisphere in MNI space (Fig. 1C). No anatomical abnormalities affecting STG regions were identified.

**Figure 1:**
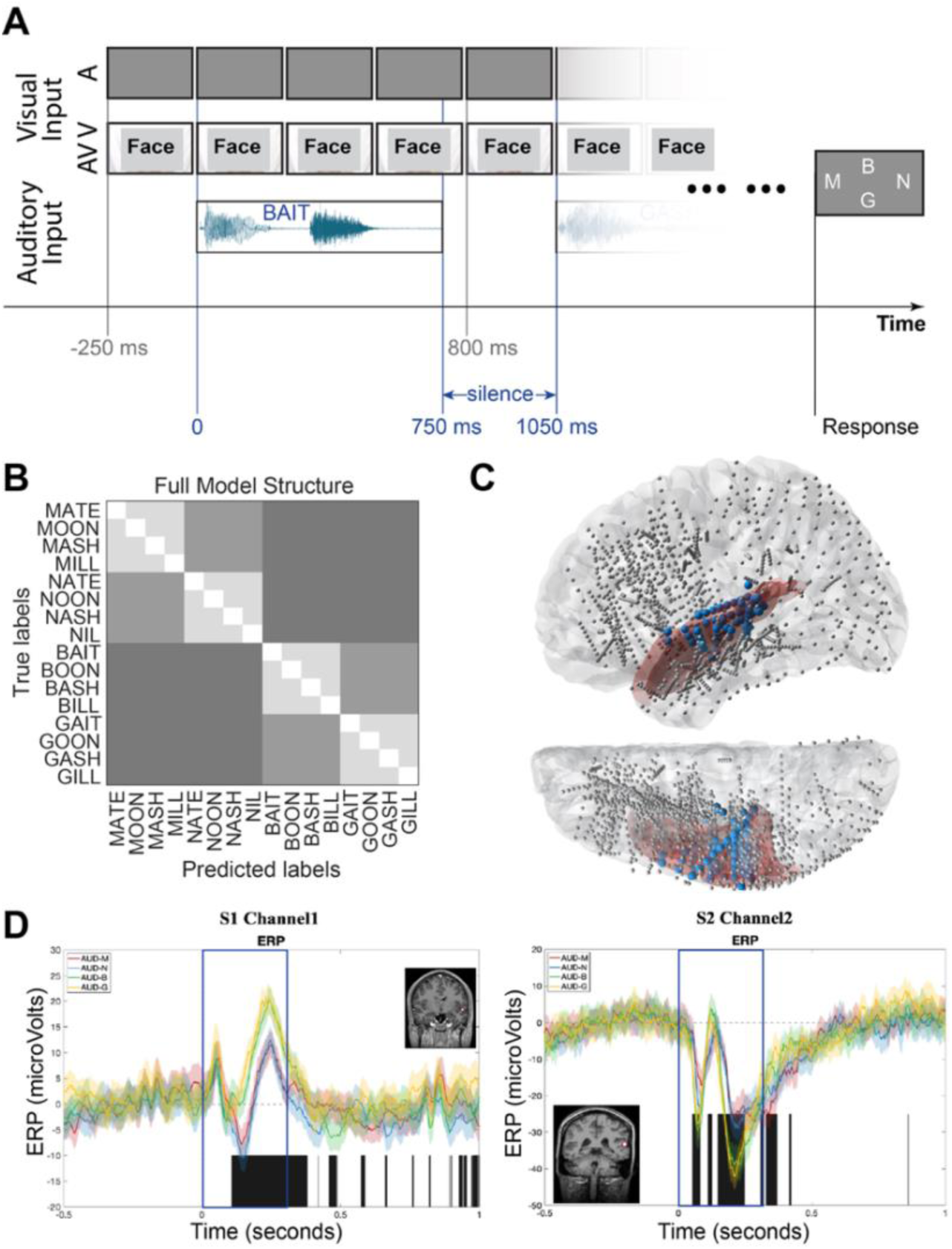
Task Schematic and iEEG Data Overview. **(A)** Trial structure. Participants heard auditory-only (A) or audiovisual (AV) word lists and identified the initial consonant of the final word. In AV trials, visual speech began 250 ms before auditory onset. **(B)** Hierarchical decoding model. The X-axis shows labels predicted by the SVM classifiers, and the Y-axis shows the true labels. Diagonal cells indicate correct word classification; off-diagonal shading denotes errors that nevertheless preserved phoneme-or phonetic-feature-level information or were incorrect at all levels. **(C)** Electrode coverage projected onto the left hemisphere in MNI space. Blue electrodes were included in the analysis, gray electrodes were excluded, and red shading highlights superior temporal gyrus (STG) grey and white matter. Electrodes from patients with bilateral coverage were projected onto the left hemisphere for visualization. **(D)** Event-related potentials (ERPs) from two representative left STG electrodes for words beginning with /b/, /g/, /m/, or /n/. Responses show phonetic-feature organization, with similar responses for nasal (/m/, /n/) and plosive (/b/, /g/) consonants. Blue boxes indicate the 0–250 ms window used for SVM classification; anatomical insets show electrode locations.

### Method details

#### Experimental design

Participants were tested at their bedside in the Epilepsy Monitoring Units using a laptop running Psychtoolbox (Brainard, 1997; Kleiner et al., 2007). The speech stimuli consisted of word lists with varying lengths, ranging from 4 to 16 words to prevent participants from attending to only the end of the list. Each list was generated by randomly selecting words from a pool of 16 monosyllabic words (for experimental setup, see Fig. 1A; for the word set, see Table S1). These words were constructed from combinations of four onset consonants (/*b*/, /*g*/, /*m*/, /*n*/) and four rimes (vowel nucleus and coda consonants; /*ey-t*/, /*uw-n*/, /*ae-sh*/, /*ih-l*/). The four onset consonants were evenly divided between plosive (/*b*/, /*g*/), and nasal (/*m*/, /*n*/) manners of articulation.

Stimuli were video recorded from a female speaker (frame rate: 59.94 fps). For each word, a single exemplar utterance was selected for presentation in three conditions: unimodal auditory (A; speech linearly mixed with pink noise, with speech and noise waveform amplitudes scaled at a 70:30 ratio, presented with a static grey screen), unimodal visual (V; video of the speaker’s face accompanied by auditory pink noise), and multimodal audiovisual (AV; synchronized auditory and visual speech, with visual onset preceding auditory onset by 250 ms). In the A and AV conditions, time 0 was defined as auditory onset, ensuring a consistent temporal reference for auditory processing across conditions. In the V condition, time 0 was instead defined as visual onset to better align neural activity associated with visual speech processing. Word durations ranged 576– 1054 ms, with onset consonants ranging 70–196 ms and rimes ranging 434–937 ms. Audio and video files were trimmed or padded to be 750 ms. Within each word list, successive words were separated by a 300-ms inter-word interval (ISI) measured from the auditory offset of one word to the auditory onset of the next. During the ISI, the final frame of the visual stimulus remained on screen until the visual onset of the subsequent word.

Participants were instructed to identify the initial consonant of the final word in each word list by button press with their dominant hand (four-alternative forced choice). Response time limits were adaptive, initially set to 2.5 s and adjusted in 250 ms steps based on performance, with bounds of 1.5 and 4 s. For each patient, trials from all three conditions were fully randomized. As testing duration varied across patients due to clinical and fatigue-related constraints, the number of repetitions differed across participants and conditions. Each word was presented at least eight times per condition (8–35 repetitions in A, and 8–32 repetitions in AV). 11 of the 12 patients completed all three conditions. One patient completed an earlier version of the task that only contained A and AV conditions.

#### Data acquisition and preprocessing

IEEG data were acquired at 4096 Hz with the Natus Quantum Amplifier (7 patients; University of Michigan Hospital, resampled to 1024 Hz) or 1000 Hz with the Nihon Kohden Amplifier (5 patients; Henry Ford Hospital). Surface reconstruction, electrode registration, and projection to Montreal Neurological Institute (MNI) coordinates were performed by aligning each patient’s preoperative T1 MRI to their postoperative CT using FreeSurfer and custom MATLAB scripts (Brang et al., 2016).

Drift was removed from each electrode by fitting and subtracting a third-order polynomial, followed by high-pass filtering at 0.1 Hz. Notch filtering at 60 Hz was applied to remove power line noise. No additional low-pass or band-pass filtering was applied, so the full broadband signal from 0.1 Hz to the Nyquist frequency (UM: 512 Hz; HF: 500 Hz) was retained for ERP computation. High-gamma power (HGp; 70–150 Hz) was extracted using wavelet decomposition for complementary analyses reported in the **Supplementary Materials**. Excessively noisy electrodes were identified and excluded if their signal variance deviated by more than 5 standard deviations in either direction from the mean across all electrodes. The remaining channels were manually inspected and channels with residual signal artifacts were removed. To minimize the influence of volume conduction on localized neural responses, bipolar re-referencing was applied. IEEG signals from adjacent electrodes were subtracted and assigned to a linearly interpolated spatial coordinate. Bipolar electrodes were included in analyses only if their nearest FreeSurfer volumetric anatomical label (projected onto the individual patient’s MRI) corresponded to STG (Desikan et al., 2006) and the signal from the electrode exhibited speech responsiveness. Speech responsiveness was defined as both a significant ERP response within 250 ms of sound onset (*i.e.*, at least one non-zero ERP time point; *p_adj* < 0.05, FDR-corrected) and significant decoding performance when unimodal auditory and multimodal audiovisual trials were pooled (*p* < 0.001; for details, see *Support Vector Machine Decoding*). All electrode screening steps were completed in patients’ individual anatomical space, and electrode coordinates were projected into MNI space for group-level visualization only. For the A and AV analyses, 158 electrodes met the anatomical criterion, of which 87 also passed the functional selection and were retained for subsequent analyses. For the V analyses, 142 electrodes met the anatomical criterion, with 81 retained after functional selection. The number of analyzed bipolar electrodes per patient ranged from 1 to 15 contacts. For A and AV analyses, patients had a mean of 7.25 analyzed contacts (*std* = 4.63; *n* = 12); for V analyses, patients had a mean of 6.75 analyzed contacts (*std* = 5.08; *n* = 11). Across patients, the final A and AV electrode set comprised 82 depth and 5 subdural electrodes, whereas the V electrode set comprised 76 depth and 5 subdural electrodes (see Fig. 1C).

### Quantification and statistical analysis

#### ERP pattern inspection

Although high-gamma power (HGp) is a commonly used index of local cortical processing in iEEG research, audiovisual speech effects in STG are often expressed in lower-frequency neural activity. In particular, visual speech has been shown to provide lateral or feedback inputs into auditory cortex through phase-resetting and cross-modal temporal alignment of low-frequency oscillations (Bastos et al., 2015; Luo et al., 2010; Mégevand et al., 2020; Schroeder et al., 2008), and recent iEEG studies have shown that pre-articulatory visual speech cues modulate low-frequency activity in STG (Karthik et al., 2021, 2024). ERPs were therefore selected as the primary measure because they are particularly sensitive to these low-frequency audiovisual processes. Analyses using HGp were performed for complementary inquiry (see Supplementary Materials).

The bipolar-referenced time series were segmented into 1.5-second epochs around the onset of the word ([-500, 1000 ms], with 0 marking auditory onset of the word). The epoch window was set to capture both preparatory speech-related movements preceding auditory onset and sustained post-onset neural responses during word processing. Epochs were averaged to compute four ERPs within each condition, corresponding to the four initial consonants of the word, and tested for significance against the pre-stimulus baseline period ([−300, −250 ms]). The baseline window was selected to occur after the auditory offset of the previous word and before the visual onset of articulatory movements in the AV condition, minimizing contamination from anticipatory speech-related activity. Fig. 1D shows ERP groupings in the unimodal auditory condition from 2 sample electrodes. All four ERP waveforms peaked at a similar latency, approximately 200 ms after word onset. Notably, the ERPs elicited by the two nasals (/*m*/ and /*n*/) and the two plosives (/*b*/ and /*g*/) closely tracked each other within their respective groups in both timing and waveform morphology.

#### Support Vector Machine decoding

##### Electrode-level word decoding

Support Vector Machine (SVM) classifiers were trained to decode the 16 words from single-trial ERP patterns for each patient. For each trial, the feature vector consisted of ERP amplitudes sampled at each time point between 0 and 250 ms after word onset from a single electrode. The decoding window was selected based on a previous implementation of the same task paradigm using a different set of monosyllabic words, which showed robust speech-related ERP responses within this time range (Karthik et al., 2024). This resulted in N-dimensional feature vectors per trial, where N corresponded to the number of sampled time points within the decoding window (*N* = 256 for UM patients and *N* = 250 for HF patients). Classification was performed at the electrode level using an 8-fold multi-class linear SVM classifier (MATLAB function ‘fitcecoc’). Multiclass classification was implemented using the one-vs-one approach, where a separate binary SVM classifier was trained to distinguish each pair of classes. With 16 classes, this approach yields a total of 120 binary classifiers. Cross-validation was stratified so that each fold contained a balanced proportion of each word label.

Classification accuracy was computed as the k-fold mean proportion of correctly predicted labels across all held-out trials. Accuracy was first calculated for each electrode and then averaged across electrodes within each patient for group-level analysis. With 16 classes per condition, the theoretical chance level was 6.25%. For each electrode, statistical significance of classification accuracy was assessed using a one-tailed binomial test (MATLAB function ‘binocdf’), comparing observed accuracy to chance. The minimum accuracy required for statistical significance depended on the number of trials and was determined individually for each electrode.

In addition to overall classification accuracy, we further evaluated decoding performance with confusion matrices and classifier confidence. Classifier confidence was quantified using the negated average binary classification loss (‘NegLoss’) returned by the MATLAB function ‘kfoldPredict’, where higher values indicated smaller error magnitude and therefore higher classifier confidence. Trial-wise NegLoss values were converted by row-wise softmax into probability-like distributions across the 16 classes, removing any additive offset shared across classes. For each electrode, these per-trial distributions were then averaged across all trials of each word to form a 16-by-16 continuous error-magnitude matrix, paralleling the structure of the discrete confusion matrix. Both confidence matrices and confusion matrices were generated for each electrode and then averaged within patients for group-level analysis.

##### Electrode-level rime decoding

To test whether audiovisual enhancement extended to rime information beyond onset consonants, additional SVM classifiers were trained to decode rime labels from ERP patterns. The decoding procedure was identical to that used for word-label decoding, except that the analysis window was time-locked to the offset of the onset consonants, with window length defined as the median rime length across all 16 words (702 ms). Phoneme timing was estimated using the Montreal Forced Aligner (McAuliffe et al., 2017), then manually verified and corrected. We first performed a 16-class rime decoding analysis in which classifiers were trained to predict the rime label associated with each of the 16 words. To further dissociate rime representations from whole-word identity, an additional 4-class SVM analysis was conducted in which separate classifiers were trained to discriminate among the four rimes regardless of the preceding consonants. Statistical significance for rime decoding performance was assessed using one-tailed binomial tests following the same procedure used for word-label decoding, except that theoretical chance performance was 6.25% for the 16-class classifiers and 25% for the 4-class classifiers.

##### Patient-level time-resolved decoding

To examine the temporal dynamics of decoding performance, time-resolved SVM analyses were additionally performed from −500 to 1000 ms relative to auditory onset. For these analyses, iEEG data were resampled to 500 Hz, yielding decoding estimates every 2 ms. For each patient and time point, the feature vector consisted of ERP amplitudes across all analyzed electrodes. Classifiers were trained and tested independently at each time point, producing patient-level time-resolved decoding accuracy curves for each condition. Decoding accuracies were then smoothed within each patient using a 100 ms (i.e., 50 time points) moving average implemented with the MATLAB function ‘movmean’. Statistical significance against chance performance at each time point was assessed using one-sample t-tests across patients and corrected for multiple comparisons using false discovery rate (FDR) correction. Successful decoding onset was defined as the first run of at least 20 ms in which group accuracy exceeded empirical chance (one-sided t-test, FDR-corrected across conditions and time points, *p* < 0.05). To compare successful decoding latencies between AV and A conditions, we used a jackknife-based procedure (Kiesel et al., 2008; Miller et al., 1998), estimating onsets from leave-one-out averages across n patients and evaluating the AV–A latency difference with a jackknife-corrected paired t-test against n − 1 degrees of freedom.

#### Hierarchical model for information processing

To answer the question of which levels of linguistic representation are susceptible to visual speech, we designed a hierarchical model to interpret patterns in the group-level confusion matrix and confidence matrix (see Fig. 1B for model structure). Rather than training separate classifiers for word-, phoneme-, and feature-level representations, we used a single word-identification model and examined the structure of its classification errors. This approach mirrors the behavioral task completed by participants and enables characterization of how different linguistic representations contribute to word-level decisions, thus providing a closer analogue to human speech perception. The diagonal elements of each matrix represent trials in which the word label was decoded correctly (word-level accuracy), while off-diagonal cells were categorized into three types to capture different levels of decoding accuracy: (1) phoneme-level accuracy, where the initial consonant was correctly decoded but the remainder of the word label was incorrect; (2) feature-level accuracy, where the initial consonant was misclassified as another consonant sharing the same manner of articulation (e.g., a word starting with ‘m’ classified as one starting with ‘n’); and (3) other errors, where neither the initial consonant nor its manner of articulation matched the true label.

This exclusive hierarchical structure enabled dissociation of decoding performance at the word, phoneme, and feature levels. Because the word set crossed four onset consonants with four rimes, each representational level was estimated from a different number of confusion-matrix cells: 16 cells contributed to the word level (the matrix diagonal), 48 to the phoneme level (correct onset, incorrect rime), and 64 to the feature level (correct shared manner of articulation, incorrect onset and rime). To determine whether audiovisual enhancement differed statistically across representational levels, audiovisual accuracy gains (AV − A) were first analyzed using a linear mixed-effects model with representational level as a fixed effect and subject as a random effect. Follow-up Wilcoxon signed-rank tests were then conducted separately at each of the four representational levels for both the confusion and confidence matrices to characterize the effect of visual speech on decoding accuracy and classifier error-magnitude within each level.

## Results

Since only the final word of each word list required a behavioral response, behavioral data were available for 1,022 of 10,285 iEEG epochs (9.9%). Participants remained engaged throughout the task, responding on the majority of trials across conditions (response rates: A: *M* = 93.9%, *std =* 6.0%; AV: *M* = 94.9%, *std =* 4.8%; V: *M* = 82.7%, *std =* 23.7%). Identification accuracy was high when auditory speech was present (A: *M* = 87.6%, *std =* 13.4%; AV: *M* = 90.5%, *std =* 15.2%) but substantially lower in the visual-only condition (*M* = 42.0%, *std =* 15.4%). Performance in the unimodal visual condition remained significantly above chance (0.25) (Wilcoxon signed-rank, *W*(12) = 75, *p* = 0.001, *r* = 0.923), indicating that participants were able to extract some speech information from visual input alone. Although AV accuracy was numerically higher than A accuracy, this difference did not reach statistical significance (*W*(13) = 53, *p* = 0.151, *r* = 0.359).

Speech-responsive electrodes in STG were selected for group-level analysis using an orthogonal channel selection procedure. Meaningful channels were identified as those with overall decoding accuracy significantly above chance, calculated across both unimodal auditory (A) and audiovisual (AV) trials combined (*p* < 0.001). Subsequent statistical comparisons of decoding accuracy between AV and A conditions were performed exclusively on this pre-selected set of channels (See Fig. 1C for the distribution of all selected channels).

We first tested whether decoding accuracy for word identity was higher in the AV condition compared to the A condition (Fig. 2A). Overall, electrode-level SVM decoding accuracy was 0.169 (*std =* 0.069) for AV, 0.158 (*std* = 0.068) for A, and 0.068 (*std* = 0.009) for V. Wilcoxon signed-rank test between AV and A revealed that AV accuracy was significantly higher than A (*W*(12) = 67, *p =* 0.013, *r* = 0.634).

**Figure 2:**
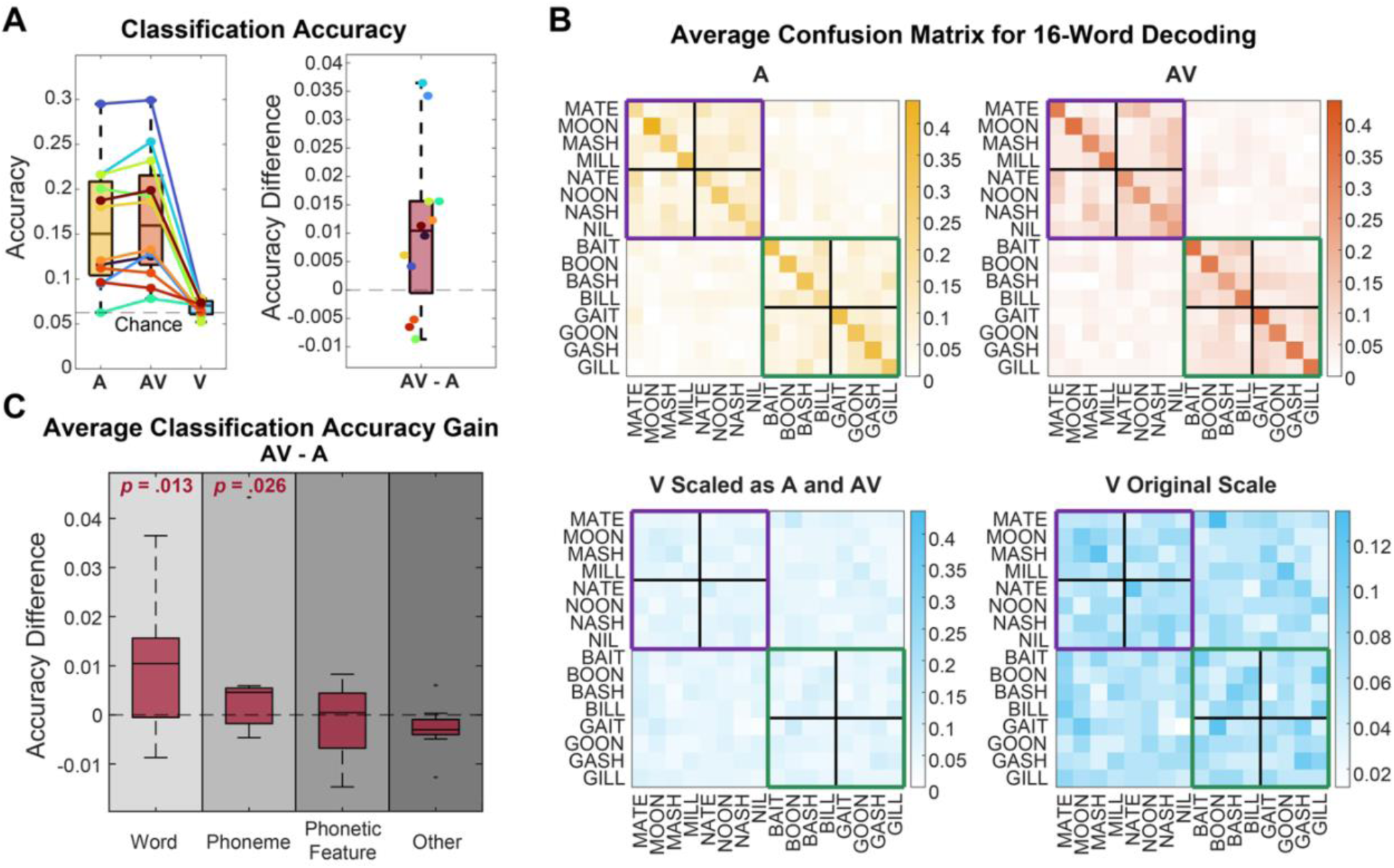
Visual speech enhances word-and phoneme-level decoding accuracy. **(A)** SVM decoding accuracy for 16-word classification across 12 patients in the auditory-only (A), audiovisual (AV), and visual-only (V) conditions. Left, mean accuracy in each condition; each colored dot represents one patient. Right, AV – A accuracy difference. Dashed lines indicate chance (0.0625) and zero difference, respectively. AV showed significantly higher decoding accuracy than A. **(B)** Group-averaged confusion matrices for A, AV, and V. The V condition is shown both rescaled to match the A/AV range and on its original scale. Axes show true versus predicted labels. Black 4x4 boxes mark correct initial-consonant (phoneme-level) decoding; colored boxes mark correct phonetic-feature decoding (purple, nasal; green, plosive). **(C)** AV – A accuracy differences at each hierarchical level (see Fig. 1B). Statistically significant differences between AV and A were observed for both word-and phoneme-level decoding.

To further determine which level of information was affected by congruent visual speech input, we used confusion matrices to decompose classifier performance in addition to examining correct responses. Even when word identity was misclassified, the predicted word could still share the target’s phoneme or phonetic features, allowing us to quantify information retained at each level (Fig. 2B-C). Audiovisual gain differed significantly across representational levels, according to the linear mixed-effects model with representational level as a fixed effect and subject as a random effect, *F*(3, 44) = 4.420, *p* = 0.008, *R^2^_marginal_* = 0.216, *R^2^_conditional_* = 0.216. Wilcoxon signed-rank tests were then conducted at each of the four levels. A significant enhancement in the AV condition was observed at the word level (A: 0.158, AV: 0.169; *W(*12*) =* 67, *p* = 0.013, *r* = 0.634) and the phoneme level (A: 0.075, AV: 0.080; *W*(12) = 64, *p* = 0.026, *r* = 0.566). No significant differences were found at the feature level (A: 0.068, AV: 0.067; *W*(12) = 35, *p* = 0.633, *r* = −0.091) or for other types of error (A: 0.043, AV: 0.040; *W*(12) = 12, *p* = 0.987, *r* = −0.611). Notably, the absence of an audiovisual effect at the feature level cannot be attributed to limited measurement precision at that level. The phonetic category was estimated from more confusion-matrix cells than either the phoneme or word level (64 versus 48 and 16 cells, respectively), so any genuine audiovisual modulation of phonetic-feature representations would have been estimated with comparable or greater statistical precision than the effects detected at the word and phoneme levels.

Classification accuracy provides a thresholded readout of representational changes, registering an effect only when sharpening is sufficient to convert a misclassification into a correct one. More gradual improvements in representational separability may therefore remain undetected if neural representations move toward the correct category but fail to cross a decision boundary and change the final classification outcome. To test for such sub-threshold effects, we next examined classifier confidence, quantified by the distance from the decision boundaries (Fig. 3). As with the confusion matrix analysis, the linear mixed-effect model revealed a significant effect of representational level on audiovisual gain, *F*(3, 44) = 3.286, *p* = 0.029, *R^2^_marginal_* = 0.170, *R^2^_conditional_* = 0.170, and follow-up Wilcoxon signed-rank tests were conducted for each of the four levels. In this case, a significant enhancement in the AV condition was observed only at the phoneme level (A: 0.066, AV: 0.067; *W(*12*) =* 71, *p* = 0.005, *r* = 0.725). No significant differences were found at the word level (A: 0.074, AV: 0.075; *W(*12*) =* 60, *p* = 0.055, *r* = 0.476), feature level (A: 0.066, AV: 0.065; *W(*12*) =* 38, *p* = 0.545, *r* = −0.023), or for other error types (A: 0.058, AV: 0.043; *W(*12*) =* 22, *p* = 0.912, *r* = −0.385). For completeness and comparison to auditory iEEG work, decoding and hierarchical analyses were also performed using HGp (70–150 Hz) with the same pipeline. HGp-based analyses showed a qualitatively similar but weaker overall pattern compared with ERP-based analyses (Fig. S1).

**Figure 3:**
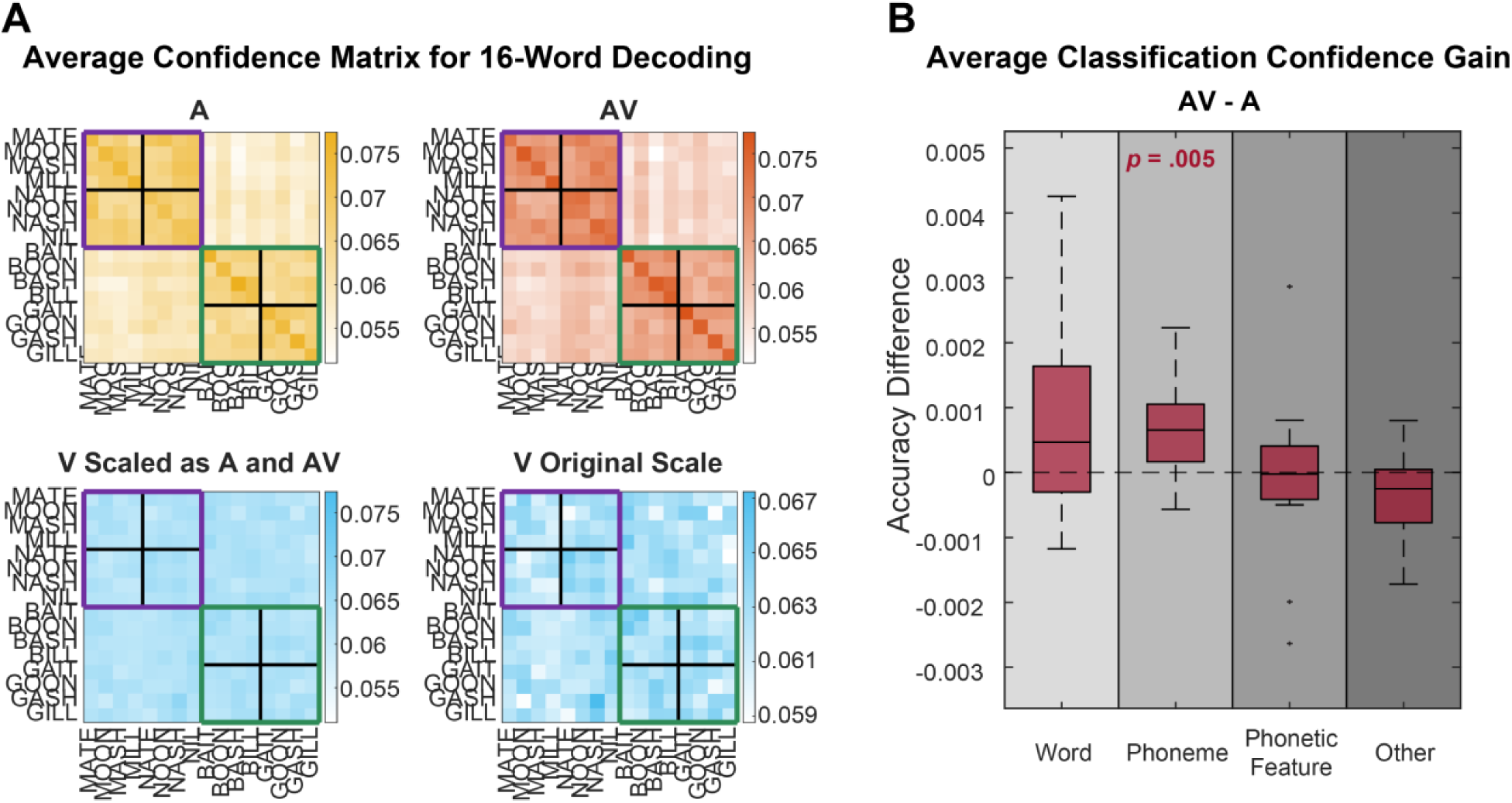
Visual speech enhances phoneme-level confidence. **(A)** Group-averaged confidence matrices for unimodal auditory (A), audiovisual (AV), and unimodal visual (V) conditions. Classifier confidence was derived from normalized inverse classification loss, with higher values indicating lower classification error and a better match between the neural activity pattern and a given class. **(B)** Statistical comparison of hierarchical model structure. The figure setup is identical to that in Fig. 2B-C, except that the values plotted here reflect classifier confidence rather than classification accuracy. Classifier confidence was quantified as being inversely related to the distance from the decision boundaries (See Methods). Statistically significant differences between AV and A were only observed for phoneme-level decoding.

Given that audiovisual enhancement was observed for word-and phoneme-level decoding accuracy, as well as phoneme-level confidence, but not for phonetic-feature representations, we hypothesized that strengthened categorical phoneme representations may contribute to the observed improvements in word-level classification. To test this hypothesis, we fit a linear mixed-effects model predicting word decoding gain from increased phoneme confidence while accounting for variability across patients. There was a non-significant positive relationship between increased phoneme confidence and word decoding gain (*b =* 2.579, *SE* = 2.14, *t*(85) = 1.205, *p* = 0.232, 95% CI [-1.677, 6.834]). Although this analysis did not provide direct statistical support, the positive coefficient was consistent with the proposed link between clarified phoneme-level representations and improved word-level decoding.

To further characterize the temporal dynamics of audiovisual enhancement, and to test whether AV benefits extended beyond onset consonants to facilitate decoding of subsequent rime information, patient-level time-resolved decoding analyses were performed at 2 ms intervals (Fig. 4A). Above-chance word-level decoding emerged earlier in the AV condition than in the A condition, starting at −2 ms and 32 ms relative to auditory onset, respectively. A jackknife-based latency comparison confirmed the earlier onset for AV than A decoding, mean AV–A difference = −35 ms, *t*(11) = −2.124, *p* = 0.029. Decoding remained elevated throughout much of the post-auditory-onset interval. Peak decoding accuracy was also higher in the AV (0.168) condition relative to the A (0.164), *W(*12*) =* 67, *p* = 0.013, *r* = 0.634. In contrast, decoding accuracy in V emerged later (118 ms) and remained comparatively modest throughout the trial (peak accuracy = 0.076).

**Figure 4:**
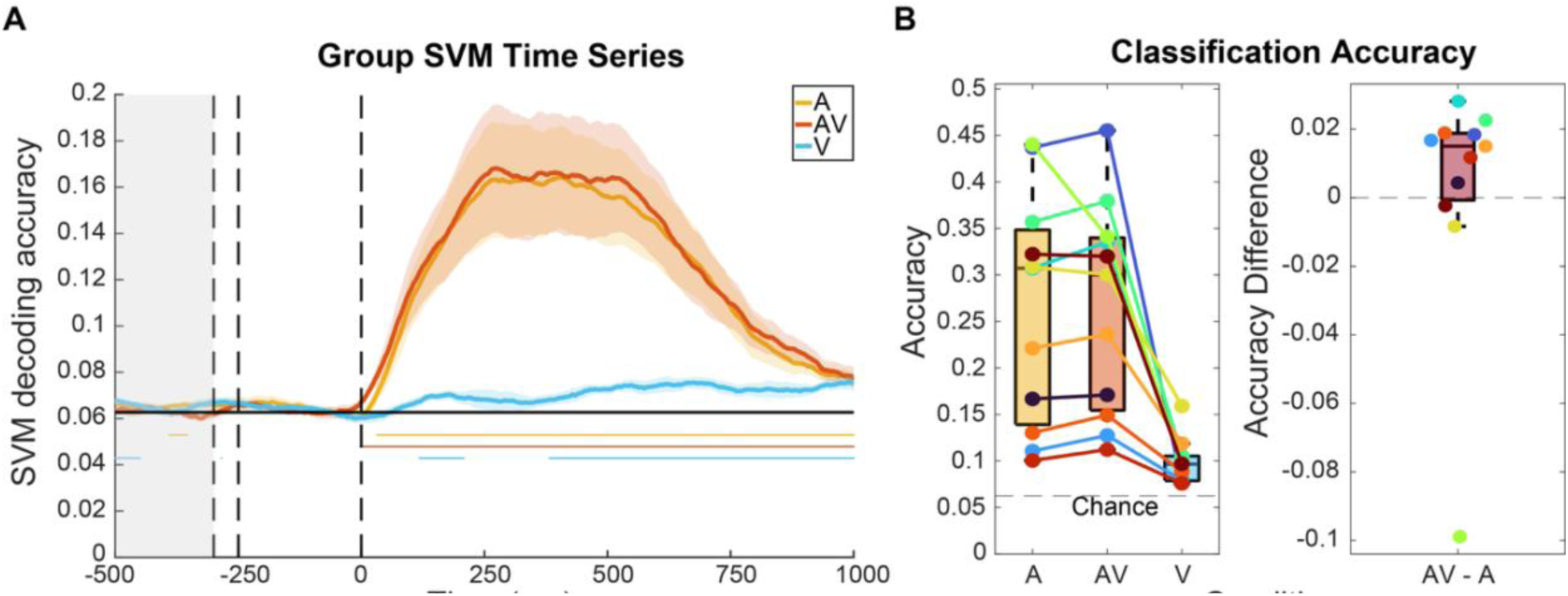
Visual speech accelerates word decoding with limited rime enhancement. **(A)** Group-averaged time-resolved SVM decoding accuracy for word labels in the auditory-only (A), audiovisual (AV), and visual-only (V) conditions across 12 patients, plotted relative to auditory onset (0 ms). Additional dashed lines indicate visual onset of the current word (−250 ms) and auditory offset of the preceding word (−300ms, grey shaded region). Colored shaded regions represent ±1 SEM across participants; colored horizontal bars indicate time points significantly above chance, *p* < 0.05, FDR-corrected). AV showed earlier decoding onset and higher peak accuracy than A, whereas V decoding emerged later and remained weaker. **(B)** Electrode-level SVM decoding accuracy for rime labels in the A, AV, and V conditions across 11 patients. One patient was excluded because no bipolar contacts met the speech-responsive criterion (significant decoding for A and AV combined, p < .001). The figure format is the same as in Fig. 2A. AV rime decoding was numerically higher than A but did not differ significantly.

Given that time-resolved decoding accuracy remained significantly above chance throughout the trial in both the A and AV conditions, we next examined whether audiovisual enhancement was also present for rime-level information (Fig. 4B). Electrode-level 16-class SVM decoding accuracy for rimes was numerically higher in AV (0.266) than that for A (0.263) at the group level, although this effect did not reach statistical significance (*W*(11) = 51, *p* = 0.062, *r* = 0.483). To determine whether this trend toward an AV advantage reflected facilitated rime processing or instead was driven by improved whole-word decoding, we conducted an additional analysis in which classifiers discriminated among the 4 rimes irrespective of the preceding consonants. Averaged patient-level time-resolved decoding performance was significantly above chance in both the A and AV conditions (Fig. 5A; peak decoding accuracy in A: 0.407, AV: 0.410, V: 0.283; significance latency relative to auditory onset: A = 76 ms, AV = 78 ms, V = 472 ms). However, no evidence for an audiovisual advantage was observed in decoding onset latency (jackknife-based comparison: mean AV-A difference = 44 ms, *t*(11) = 0.134, *p* = 0.552), peak decoding accuracy (*W*(12) = 44, *p* = 0.367, *r* = 0.113), or in the electrode-level decoding accuracy (Fig. 5B; *W*(11) = 33, *p* = 0.517, *r* = 0.0). Together, these findings suggest that the AV-related improvement observed in the 16-class rime analysis was more likely attributable to enhanced whole-word representations than to rime information itself.

**Figure 5:**
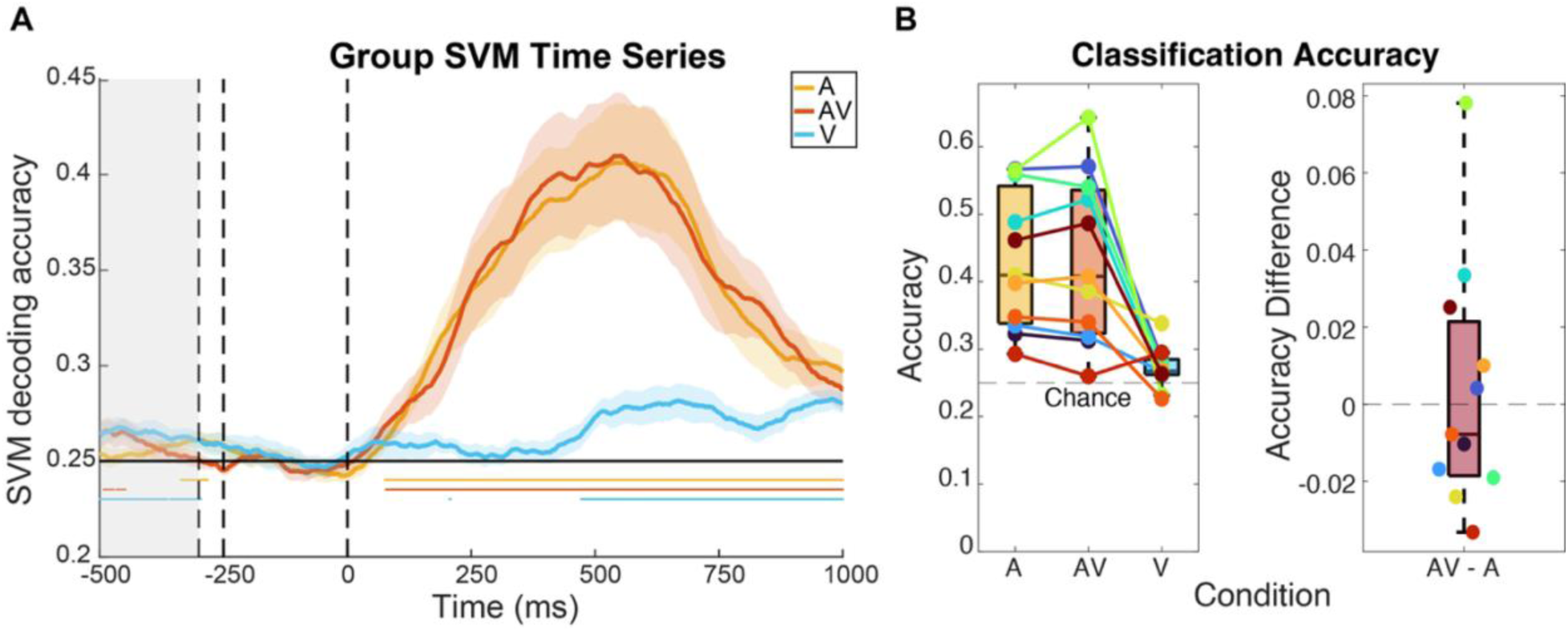
Rime enhancement reflects word-level audiovisual advantage. **(A)** Group-averaged time-resolved 4-class SVM decoding accuracy for rime identity in the auditory-only (A), audiovisual (AV), and visual-only (V) conditions. The figure format is the same as in Fig. 4A. **(B)** Electrode-level 4-class SVM decoding accuracy for rime identity across the 11 included patients in the A, AV, and V conditions. The figure format is the same as in Fig. 4B, except that decoding was performed across the four rimes irrespective of onset consonant. Although decoding in A and AV remained above chance, no significant audiovisual advantage was observed in either the time-resolved or electrode-level analysis.

## Discussion

Congruent visual input facilitates speech perception, but this behavioral enhancement can arise through multiple mechanisms, including changes in temporal processing (Cao et al., 2024; McGrath & Summerfield, 1985; Mégevand et al., 2020), attention allocation (ten Oever et al., 2014; Zion Golumbic et al., 2012; Zion Golumbic et al., 2013), cue weighting (Shahin et al., 2018), speech parsing (Chandrasekaran et al., 2009), and potential integration of spectral (Bröhl et al., 2022; Plass et al., 2020) and visemic information (Karthik et al., 2021) derived from visual cues. In addition to vision-driven modulation of auditory responses during speech perception (Okada et al., 2013), recent evidence demonstrates that visemic identities are directly represented in the auditory cortex in a manner analogous to phonemes (Audenhaege et al., 2025; Karthik et al., 2024), suggesting a direct interaction with linguistic representations in the auditory cortex. However, it remains unclear whether such audiovisual integration alters speech representations encoded by neuronal populations in STG, and if so, at which level of linguistic representation these effects emerge. In the present study, we showed enhanced neural decoding accuracy for word identity in audiovisual relative to unimodal auditory conditions, which is consistent with classic findings from the speech perception literature and extend prior findings to neural representations of audiovisual speech. Importantly, our results suggest that the improvement in word-level decoding accuracy may arise from enhanced confidence in phoneme-level representations.

Although the increase in phoneme-level classifier confidence and the improvement in word-level decoding accuracy were observed in different metrics and at different levels of the hierarchical model, they likely reflect a common underlying change in representational structure. Classification accuracy is a thresholded measure that captures discrete representational changes only when they alter classification outcomes, whereas classifier confidence is sensitive to more subtle continuous changes in representational separability. Visual speech may therefore first augment phoneme representations, increasing confidence before affecting decoding accuracy. As phoneme representations become increasingly distinct, these changes eventually improve both phoneme-and word-level classification. Since the words in our word set were primarily distinguished by onset phoneme identity, increased phoneme separability would be expected to propagate to word-level decoding. Consistent with this interpretation, the positive coefficient in the LME analysis suggests that the phoneme-confidence and word-decoding effects may reflect successive stages of the same representational process.

The absence of any audiovisual effect at the phonetic-feature level further constrains the nature of this representational change. Importantly, this finding does not suggest that phonetic features are weakly represented in STG. In fact, the confusion and confidence matrices showed clear phonetic-feature grouping in both unimodal auditory and audiovisual conditions, consistent with prior work showing that phonetic features are among the most salient dimensions encoded in human STG (Leonard et al., 2024; Mesgarani et al., 2014). Thus, the dominant representational structure in the present task remained organized by phonetic features, as expected for an auditory-dominant speech task.

However, our central question was which level of representation is modified by congruent visual speech. From this perspective, the lack of such a feature-level effect supports a genuine representational dissociation: congruent visual speech selectively increased the discriminability of phoneme representations without corresponding enhancement of the already robust phonetic-feature organization. This supports the view that audiovisual facilitation operates on categorical, lexically relevant units rather than on the acoustic-articulatory features from which those categories are formed (Shahin et al., 2018). Intracranial recordings from pSTG directly support this notion, showing that visual speech suppresses auditory responses most for mouth-leading words, an effect attributed to inhibition of incompatible phoneme representations (Karas et al., 2019). Reduced overall responses and sharper category separation are compatible outcomes of the same competitive process. This interpretation aligns well with prior research using masking (Shahin et al., 2012) and speech degradation (Rennig & Beauchamp, 2022), suggesting that visual speech cues facilitate the recovery of higher-level linguistic information and improve intelligibility of speech in noise. Likewise, silent lip-reading studies show that visual speech processing extracts visemes, the visual analogues of phonemes, dissociably from lip movements themselves (Nidiffer et al., 2023). Taken together, these findings support the view that visual information contributes to the identification of abstract, meaning-related units or provides complementary cues to linguistic content during word processing, rather than simply recovering missing acoustic or articulatory details.

The temporal dynamics of whole-word decoding reported here showed earlier successful decoding in the audiovisual condition relative to the unimodal auditory condition. This finding supports a predictive role for visual speech. As articulatory movements typically precede the voice, visual cues constrain predictions for upcoming phonetic content, shortening auditory response latencies in proportion to visual predictiveness (van Wassenhove et al., 2005) via a fast route conveying visual predictions directly to auditory cortex (Arnal et al., 2009). The temporal window for audiovisual integration is wide and asymmetric (van Wassenhove et al., 2007), aligning with the multi-scale integration windows proposed for speech processing (Poeppel et al., 2008). Collectively, these mechanisms allow visual input to sharpen auditory representations before the acoustic signal is fully available, consistent with the earlier word-level decoding observed here. Notably, patient-level time-resolved decoding analyses showed robust rime decoding in both audiovisual and unimodal auditory conditions, indicating that sustained speech information remained decodable throughout the trial. However, significant group-level audiovisual enhancement in the electrode-level decoding analysis was observed for onset consonants, but not for rimes. These findings are consistent with previous work showing that visual speech primarily benefits the initial stages of auditory speech processing, whereas sustained speech processing relies more heavily on tracking the acoustic envelope (Cao et al., 2024). This distinction is further supported by converging intracranial evidence. Words with a visual head start show the largest visual benefit (Karas et al., 2019), and human STG distinguishes onset from sustained responses (Hamilton et al., 2018).

Additionally, we found that 16-class rime decoding, which preserved coarticulatory information between onset consonants and rimes, showed a greater numerical audiovisual advantage than the 4-class rime decoding analysis. This pattern suggests that visual speech may be particularly informative for distinguishing transitional acoustic-phonetic patterns rather than isolated phoneme categories, consistent with the inherently coarticulated nature of continuous speech (Schwartz & Savariaux, 2014; Venezia et al., 2016). Within the current experiment, the stronger audiovisual effect observed in the 16-class analysis further suggests that the apparent rime advantage was largely driven by enhanced whole-word decoding rather than improved decoding of isolated rime segments. More broadly, these findings are consistent with renewed interest in the role of coarticulation in natural speech processing (Kent & Minifie, 1977; Moreira et al., 2025).

While our decoding results suggest that congruent visual information leads to better word identity decoding through more confident phoneme categorization, the underlying neural computations responsible for this enhancement remain unclear. Given that visual information is already present in the auditory cortex before sound onset (Karthik et al., 2024), future studies could adopt an encoding perspective to directly investigate the content and timing of congruent visual speech representation. Additionally, examining the spectral profile of these effects across frequency bands may help clarify the neural mechanism by which visemic information is transformed into phoneme representations (Karthik et al., 2021; Schroeder et al., 2008).

The present study focused exclusively on STG. Future work incorporating regions implicated in audiovisual integration (e.g., pSTS: Audenhaege et al., 2025; Erickson et al., 2014; Karthik et al., 2024; Rennig & Beauchamp, 2022; Zhu & Beauchamp, 2017) and word retrieval (e.g., IFG: Yu et al., 2025) would provide a more comprehensive picture of information flow during multisensory speech perception. Specifically, the absence of an audiovisual effect at the phonetic-feature level in STG does not preclude such effects elsewhere in the speech network. We would predict stronger audiovisual enhancement of phonetic-feature representations in the pSTS, where visual articulatory information may be integrated before being converted into phoneme representations in STG (Zhang et al., 2025). Within STG, a sharp functional boundary separates anterior and posterior subregions in multisensory speech processing (Ozker et al., 2017, 2018), so audiovisual effects may also vary along this axis. Additional limitations include restricted phoneme coverage and imbalanced word frequencies, both necessary for the controlled word set. These constraints, however, do not compromise our central inquiry into how visual information influences speech perception, as identical stimuli were presented across conditions. A broader characterization of phoneme and viseme confusion patterns in English (Cutler et al., 2004; Fisher, 1968) will be necessary to draw more generalizable conclusions about how visemes and phonemes interact to support successful speech perception. Furthermore, extending the current analysis from word identification to longer temporal scales could yield a deeper understanding of the multisensory hierarchy underlying natural speech perception (Gwilliams, Marantz, et al., 2025; Heilbron et al., 2022; O’Sullivan et al., 2021).

In sum, the present study provides new evidence that congruent visual information enhances phoneme processing in STG, with increased confidence in phoneme identity leading to more accurate word perception. These findings are consistent with the hypothesis that visual information primarily supports the extraction of higher-level linguistic representations or provides supplementary cues for speech comprehension, rather than simply enhancing acoustic or articulatory information, encouraging future work toward a multisensory framework for speech perception.

## Author Contributions

DB, VW, and WS designed the study. Data were collected by VW, WS, YL, and DB. VW, WS, and DB provided guidance regarding clinical data access and preprocessing. YL, ID, and DB developed methods for data analysis and interpretation. YL wrote the initial manuscript. All authors contributed to manuscript revision and approved the final version.

## Conflict of interest statement

The authors declare no competing financial interests.

## Data/code availability

Code will be publicly available upon publication of this article. Data are available upon reasonable request.

## Acknowledgments

This study was supported by NIH grants R01DC020717, and R01NS094399. We sincerely thank the patients for generously contributing their time and effort to this research.

## Supplementary Materials

### HGp analyses

For completeness and comparison to other iEEG research, the same analyses were also ran on HGp. After minimal preprocessing steps (see Methods), HGp (70–150 Hz) was extracted using wavelet decomposition and used to train a separate set of electrode-level SVM decoders to decode word labels of the 16 word stimuli. Following the same inclusion criteria for STG restriction and speech responsiveness as in the ERP analyses, electrodes showing significant word decoding when A and AV trials were combined (*p* < .001) were retained for subsequent statistical and hierarchical analyses. For the A and AV analyses, 81 electrodes met the anatomical criterion, of which 20 also passed the functional selection and were retained for subsequent analyses. For the V analyses, 75 electrodes met the anatomical criterion, with 18 retained after functional selection. The number of analyzed bipolar electrodes per patient ranged from 1 to 6 contacts. For A and AV analyses, patients had a mean of 3.33 analyzed contacts (*std* = 1.97; *n* = 6); for V analyses, patients had a mean of 3.00 analyzed contacts (*std* = 2.37; *n* = 5). All electrodes included in HGp analyses were depth electrodes (See Fig. S1A).

Overall, electrode-level SVM decoding accuracy was 0.097 (*std =* 0.026) for AV, 0.098 (*std* = 0.012) for A, and 0.060 (*std* = 0.011) for V. Wilcoxon signed-rank test revealed no significant difference in word-label decoding accuracy between AV and A conditions (*W*(6) = 7, *p =* 0.594, *r* = −0.060). Although linear mixed-effects model showed no significant accuracy difference across representational levels, *F*(3, 20) = 1.300, *p* = 0.302, *R^2^_marginal_*= 0.140, *R^2^_conditional_* = 0.140, for completeness of analyses, we still performed Wilcoxon signed-rank tests for each of the four levels (Fig. S1C-D). A significant enhancement in the AV condition was observed only at the phoneme level (A: 0.080, AV: 0.089; *W(*6*) =* 19, *p* = 0.047, *r* = 0.728). No significant differences were found at the word level (A: 0.102, AV: 0.094; *W(*6*) =* 7, *p* = 0.594, *r* = −0.060), feature level (A: 0.075, AV: 0.073; *W(*6*) =* 3, *p* = 0.953, *r* = −0.642), or for other error types (A: 0.045, AV: 0.046; *W(*6*) =* 9, *p* = 0.656, *r* = −0.128).

Similarly, linear mixed-effects model showed no significant confidence difference across representational levels, *F*(3, 20) = 1.204, *p* = 0.334, *R^2^_marginal_*= 0.131, *R^2^_conditional_* = 0.131, and for completeness of analyses, we still performed Wilcoxon signed-rank tests for each of the four levels (Fig. S1E-F). No significant differences were observed at the word level (A: 0.074, AV: 0.073; *W(*6*) =* 15, *p* = 0.219, *r* = 0.385), phoneme level (A: 0.068, AV: 0.069; *W(*6*) =* 18, *p* = 0.078, *r* = 0.642), feature level (A: 0.068, AV: 0.067; *W(*6*) =* 12, *p* = 0.422, *r* = 0.128), or for other error types (A: 0.057, AV: 0.057; *W(*6*) =* 7, *p* = 0.781, *r* = −0.300).

**Figure S1:**
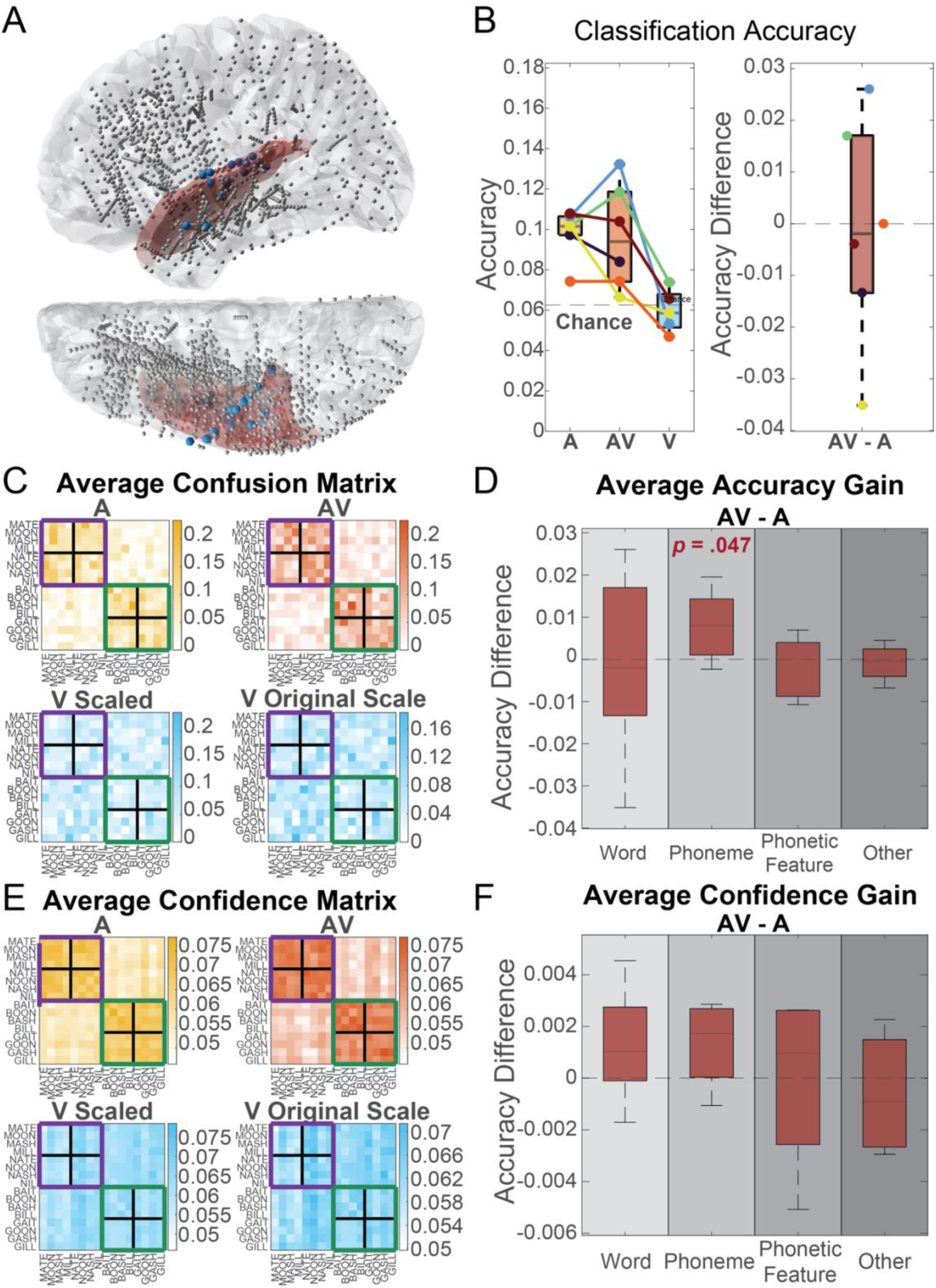
SVM Decoding for word labels and hierarchical analyses using HGp. Electrode-level SVM decoding results for 16 word labels using HGp. Six of twelve patients had at least one auditory-responsive electrode showing above-chance decoding performance when A and AV trials were combined. Panels C and D show hierarchical analyses of discrete decoding outcomes from the confusion matrix for each condition, and panels E and F show the corresponding analyses of graded representational shifts based on classifier confidence. The figure layout is identical to that of Figs. 2 and 3. HGp-based analyses showed a similar pattern to the ERP-based results, with phoneme-level, but not feature-level, representations modulated by congruent visual speech.

**Table S1:**
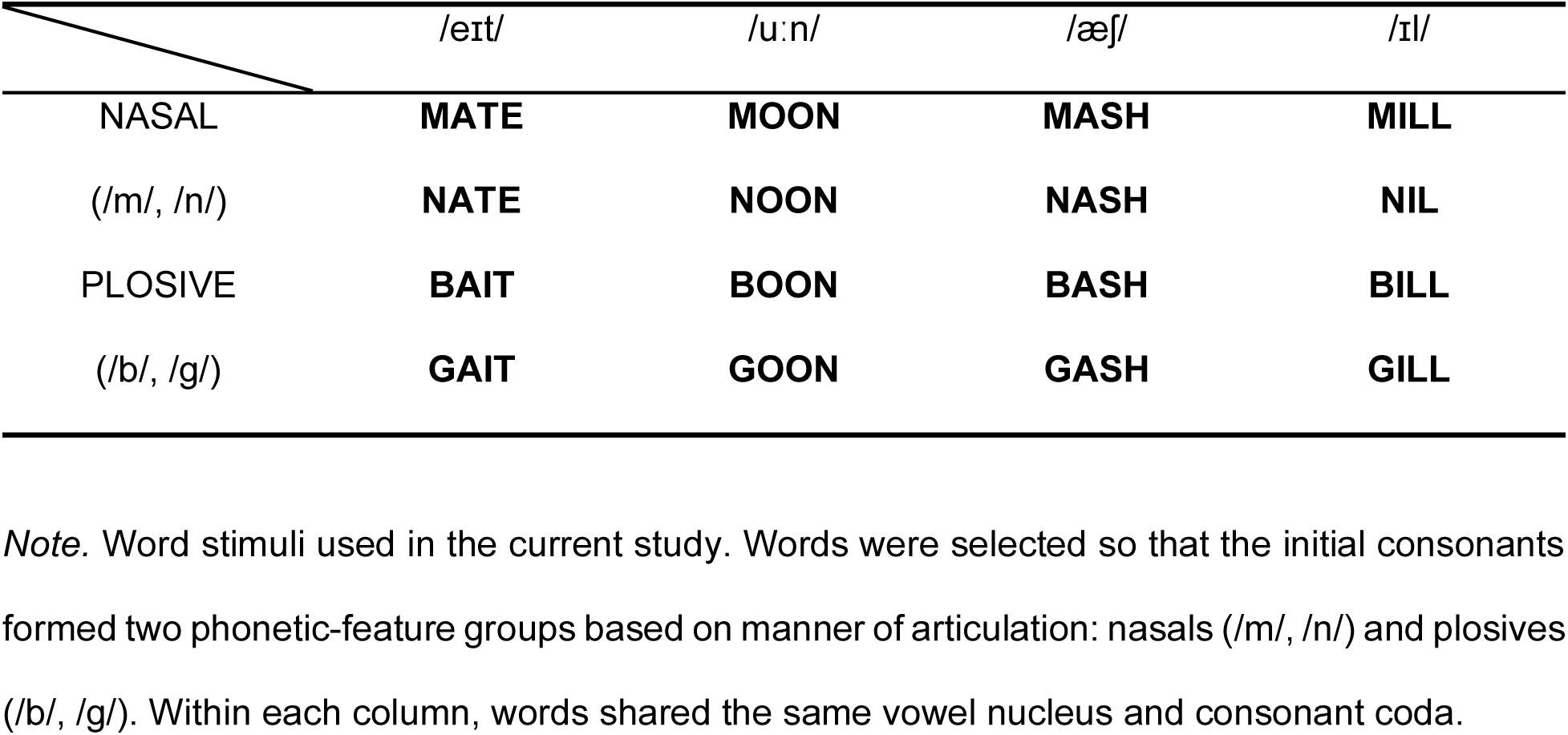
Complete word set in the study.

|  | /eɪt/ | /u:n/ | /æʃ/ | /ɪl/ |
| --- | --- | --- | --- | --- |
| NASAL | **MATE** | **MOON** | **MASH** | **MILL** |
| (/m/, /n/) | **NATE** | **NOON** | **NASH** | **NIL** |
| PLOSIVE | **BAIT** | **BOON** | **BASH** | **BILL** |
| (/b/, /g/) | **GAIT** | **GOON** | **GASH** | **GILL** |

*Note.* Word stimuli used in the current study. Words were selected so that the initial consonants formed two phonetic-feature groups based on manner of articulation: nasals (/m/, /n/) and plosives (/b/, /g/). Within each column, words shared the same vowel nucleus and consonant coda.

## Notes

### Competing Interest Statement

The authors have declared no competing interest.

